# Molecular and biophysical requirements for B cell receptor dependent enhancement of dengue virus infection

**DOI:** 10.64898/2026.01.05.697627

**Authors:** Gaby Madrigal, Chad Gebo, Benjamin D. McElvany, Sean A. Diehl, Adam T. Waickman

## Abstract

Dengue virus (DENV) is the causative agent of dengue, a mosquito-borne disease that represents a significant and growing public health burden around the world. A unique pathophysiological feature of dengue is immune-mediated enhancement, wherein preexisting immunity elicited by a primary infection can enhance the severity of a subsequent infection by a heterologous DENV serotype. A leading mechanistic explanation for this phenomenon is antibody dependent enhancement (ADE), where sub-neutralizing concentrations of DENV-specific IgG antibodies facilitate entry of DENV into FcγR expressing cells. Accordingly, this model posits that phagocytic mononuclear cells are the primary reservoir of DENV. However, multiple independent groups have shown that B cells are the largest reservoir of virally infected cells in circulation during acute dengue, representing a disconnect in our understanding of immune-mediated DENV tropism. In response to this persistent knowledge gap, our team has previously identified a novel mechanism of immune-mediated enhancement we have termed BCR-dependent enhancement (BDE) of DENV infection. In this study, we show that DENV infection of DENV-reactive B cells is highly sensitive to BCR/DENV envelope (E) protein interactions. DENV entry into this subset of B cells is dynamin-mediated and requires proximal BCR signaling. Finally, we show that DENV-reactive B cells are productively infected by live DENV, capable of supporting active viral replication and dissemination. We propose that BDE provides an additional layer of pathogen-specific immune-mediated infection risk that complements existing models of ADE and offers additional insight into potential mechanisms of DENV immunopathogenesis.

## INTRODUCTION

Dengue virus (DENV) is a prevalent arboviral pathogen that represents a significant public health burden in many regions of the world. An estimated 3.6 billion people live in regions with sustained DENV transmission, with the virus infecting an estimated 400 million individuals every year (1, 2). Transmitted via the bite of infected *Aedes* mosquitoes, DENV co-circulates as four genetically and immunologically distinct serotypes: DENV-1, -2, -3, and -4. Approximately 25% of DENV infections are thought to result in some form of symptomatic illness, ranging in severity from dengue fever - a mild flu-like presentation of infection – to life-threatening severe dengue, dengue hemorrhagic fever (DHF) or dengue shock syndrome (DSS) (3-5). Currently, there are no accurate prognostic indicators of which patients will progress to develop severe dengue. However, the risk of developing severe dengue increases in patients previously infected by a different (heterologous) serotype of dengue compared to DENV-naïve individuals (1, 6). This increased risk of severe disease following heterologous reinfection strongly suggests a mechanism of immune-mediated enhancement of DENV infection.

While the etiology of severe dengue is complex and incompletely understood, *in vitro* studies and epidemiological modeling suggest that antibody-dependent enhancement (ADE) of DENV infection may play a significant role in both the risk of DENV infection and in dengue pathogenesis (6-9). Waning titers of cross-reactive DENV-specific antibodies to sub-neutralizing levels have been posited to result in opsonization of the virion and subsequent uptake by Fcγ-receptor (FcγR) bearing cells such as monocytes, macrophages, and dendritic cells (10, 11). This has been suggested to result both in an increase in the number of infected cells early after inoculation, as well as a qualitatively altered intrinsic response to infection in these susceptible phagocytes (10, 12-14). The delicate balance between protective and enhancing levels of immunity in dengue is a hallmark of the disease and has complicated dengue control efforts for decades.

While the ADE model of dengue immunopathogenesis has been a cornerstone of the field for nearly 50 years, numerous observations strongly suggest that other immunologic mechanisms must also contribute to DENV pathology. Most notably, while *in vitro* infection studies and histological analysis of samples collected from individuals who have succumbed to severe dengue have shown a high DENV burden in FcγR expressing phagocytes (12, 15-18), *ex vivo* analysis of PBMC collected during acute DENV infection instead shows that B cells are the dominant circulating cellular reservoir of DENV (12, 19-24). This observation has been recapitulated by multiple groups - including our own - using an array of assays including flow cytometry, scRNAseq, intrathoracic inoculation of mosquitoes with purified cell populations, and qRT-PCR (12, 19-24). While B cells express an FcγR, the specific isoform of this receptor (FcγrIIb) has been shown to be incapable of facilitating ADE and instead results in anergy and apoptosis in B cells when engaged (20, 25). Accordingly, there is need for the field to reconcile these disparate observations and assumptions regarding DENV cellular tropism (12, 20, 22).

Recently, our team identified a novel DENV entry mechanism specifically within DENV-reactive B cells, a process we have termed B cell receptor-dependent enhancement (BDE). Our group, as well as others, have shown that viruses are able to utilize B cell receptor (BCR) antigen specificity in order to preferentially infect subsets of the B cell population in a highly antigen-specific fashion (26, 27). We have shown *in vitro* that B cells expressing a DENV envelope protein-specific BCR were susceptible to DENV infection, and that the frequency of DENV-infectible B cells in PBMCs increased significantly after a primary DENV infection (26). However, questions remain regarding the signaling pathways and cellular processes required for BCR-mediated infection of DENV-specific B cells.

In this study, we define the cell-intrinsic mechanistic requirements for BCR-mediated entry of DENV into DENV-specific B cells utilizing a unique set of immortalized flavivirus-specific human B cells. Using this model system, we confirm that the susceptibility of these cells is dependent both on the antigen-specificity of the BCR as well as canonical BCR signaling and endocytic pathways. In addition, we demonstrate that this antigen-specific endocytic process results in localization of DENV into late endosomes in DENV-specific B cells, an essential part of the DENV entry cycle. Finally, we confirm the ability of DENV-reactive B cells to support productive DENV replication. Overall, our results highlight the entry and signaling requirements in BDE and emphasizes how generation of DENV-specific immunologic memory may play a critical cell-intrinsic role in determining DENV susceptibility at an organismal level.

## RESULTS

### BCR antigen-specificity determines DENV susceptibility of B cells

In our prior work, we demonstrated that ectopic expression of DENV-specific BCRs rendered normally non-infectible cells highly susceptible to DENV infection (26). Furthermore, we determined that immortalized human B cell lines naturally expressing DENV-specific BCRs were uniquely and potently susceptible to DENV infection (26). To further define the mechanistic requirements for this BCR-dependent infection process, we further examined these flavivirus-reactive immortalized B cells, generated from individuals previously infected with DENV-1 and DENV-2 (7B9^DENV^ cells) or Zika virus (ZIKV, 2F3^ZIKV^ cells).

Consistent with the binding specificity of the secreted antibody produced by these cells (26), we observed robust binding of recombinant DENV-2 and DENV-4 stabilized E dimer proteins (recE) (28) to the surface of the 7B9^DENV^ cells, but not 2F3^ZIKV^ cells (**Fig. 1A, and Sup. Fig 1**). Mechanistically, if this epitope/paratope interaction contributed to the process of BDE, we hypothesized that blocking DENV-E protein binding sites on the BCR of DENV-specific B cells, would result in a reduction in the susceptibility of DENV-reactive B cells to DENV infection. To facilitate the specific quantification of productive infection – rather than antigen-binding/uptake – we utilized a DENV-pseudotyped reporter virus particle (RVP) platform. This system encodes a fluorescent reporter expressed only upon successful cytoplasmic entry and translation, allowing specific quantification of productive infection rather than surface binding alone (29, 30). Consistent with our previously published results (26), the 7B9^DENV^ cells were highly susceptible to DENV RVP infection, while the 2F3^ZIKV^ cells exhibited no RVP infection across all tested infection conditions (**Fig. 1B, and Sup. Fig. 2**).

**Figure 1.**
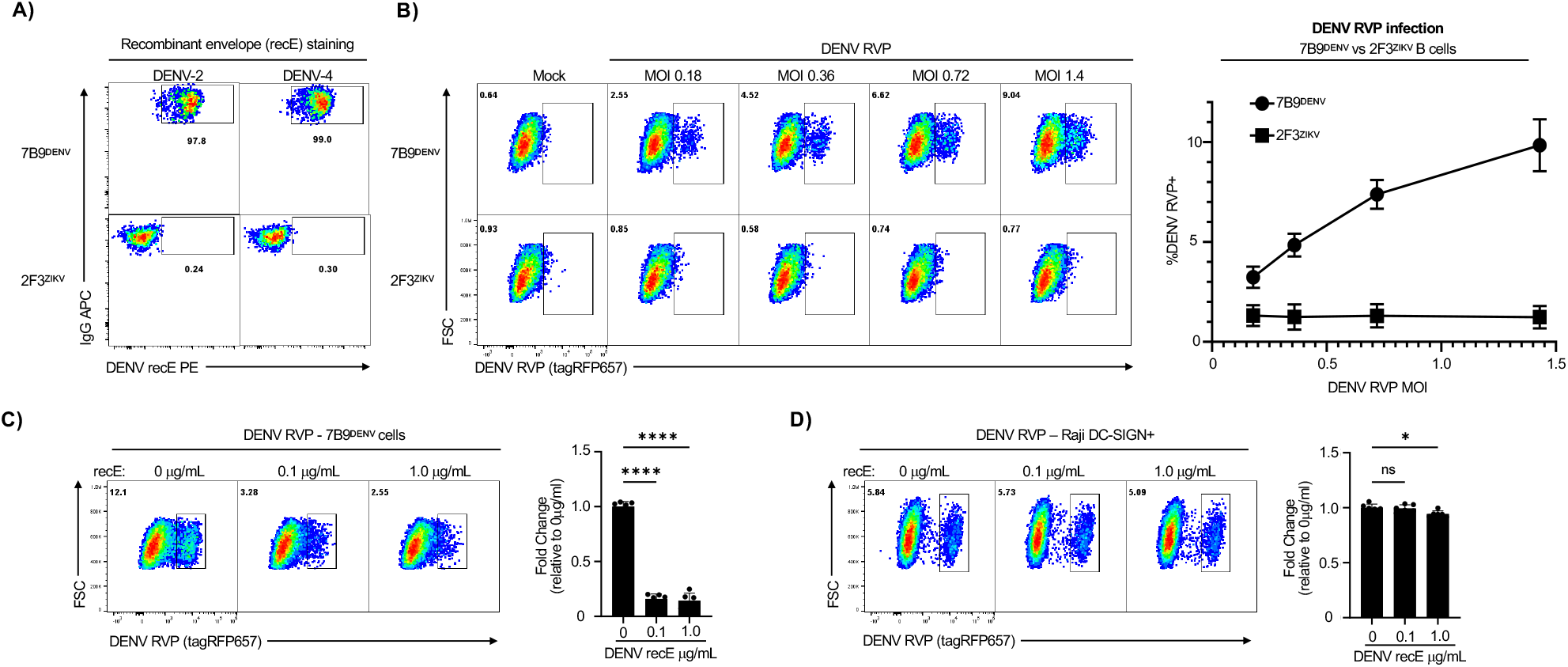
BCR-dependent enhancement in DENV-reactive B cells is highly sensitive to DENV Envelope-specific interactions. **A)** Binding of conformational recombinant E dimers (recE) from DENV-2 (left) and DENV-4 (right) to 7B9^DENV^ and 2F3^ZIKV^ immortalized B cells. **B)** Flow cytometric analysis of DENV-4 pseudotyped reporter virus particle (RVP) titration on 7B9^DENV^ and 2F3^ZIKV^ cells. Data collected 24 hours post infection. **C)** Flow cytometric analysis of DENV-4 RVP infected 7B9^DENV^ cells pre-treated with 0.1 μg/mL or 1.0 μg/mL of DENV-2 recE before addition of DENV-4 RVP. **D)** Flow cytometric analysis of Raji DC-SIGN+ cells pre-treated with 0.1 μg/mL or 1.0 μg/mL of DENV-2 recE before addition of DENV-4 RVP. Error bars +/-SEM. ns = not significant, ^*^ *P* < 0.05, ^****^ *P* < 0.0001, One-way ANOVA with Dunnett’s multiple comparisons test, with a single pooled variance. Experiments were conducted across two biological replicates in triplicate.

We next incubated 7B9^DENV^ cells with recE prior to the addition of DENV RVP. As hypothesized, we observed a dose-dependent reduction in the frequency of RVP-infected cells in the presence of recE blockade (**Fig. 1C**). To confirm this reduction in infection was due to the BCR specificity of the 7B9^DENV^ cells and not due to other antagonistic interactions, we repeated this recE blockade in a known DENV-susceptible Raji DC-SIGN+ cell line. DC-SIGN/DC-SIGNR-mediated DENV uptake, unlike BCR-mediated uptake, depends on low-affinity, high-valency interactions with E protein carbohydrates rather than specific epitope recognition (31, 32). Accordingly, we predicted– in contrast to what was observed for BCR-mediated infection – that DC-SIGN/DC-SIGNR-mediated infection would be minimally impacted by the exogenous addition of soluble recE. Consistent with this model, we only observed a modest reduction in RVP infection of DC-SIGNR expressing Raji cells (**Fig. 1D**), emphasizing the specificity and potency of BCR-mediated DENV uptake and subsequent infection.

### BCR signaling is required for BDE-mediated viral entry

Having demonstrated the role for BCR specificity in DENV infection of immortalized human B cells, we next sought to define the cell-intrinsic signaling and biophysical processes required for BDE. We posited that BCR-mediated infection of DENV-specific B cells would be dependent on canonical B cell signaling and endocytic processes including dynamin-mediated endocytosis and Bruton’s tyrosine kinase (BTK) signaling. However, we predicted that other downstream BCR signaling processes not directly involved in endocytosis – such as PI3K signaling – would be dispensable for the initial entry of DENV into the B cell, although these pathways may be involved in regulating the permissivity of the cell to DENV infection due to their role in regulating protein translation and cell cycle progression.

To address these questions, we utilized pharmacological inhibitors of endocytosis and proximal BCR signaling, specifically targeting dynamin-mediated endocytosis, Bruton’s tyrosine kinase (BTK) signaling, and phosphoinositide 3-Kinase (PI3K) signaling (**Fig. 2A**). 7B9^DENV^ cells were pre-treated for 30 minutes with the indicated inhibitors prior to the addition of the DENV RVP followed by an additional 4 hour incubation in presence of inhibitors for infection. Following extensive washing and trypsin-treatment to remove any remaining surface bound RVP, cells were incubated for an additional 20 hours and analyzed by flow cytometry to quantify the frequency of DENV RVP infected cells.

**Figure 2.**
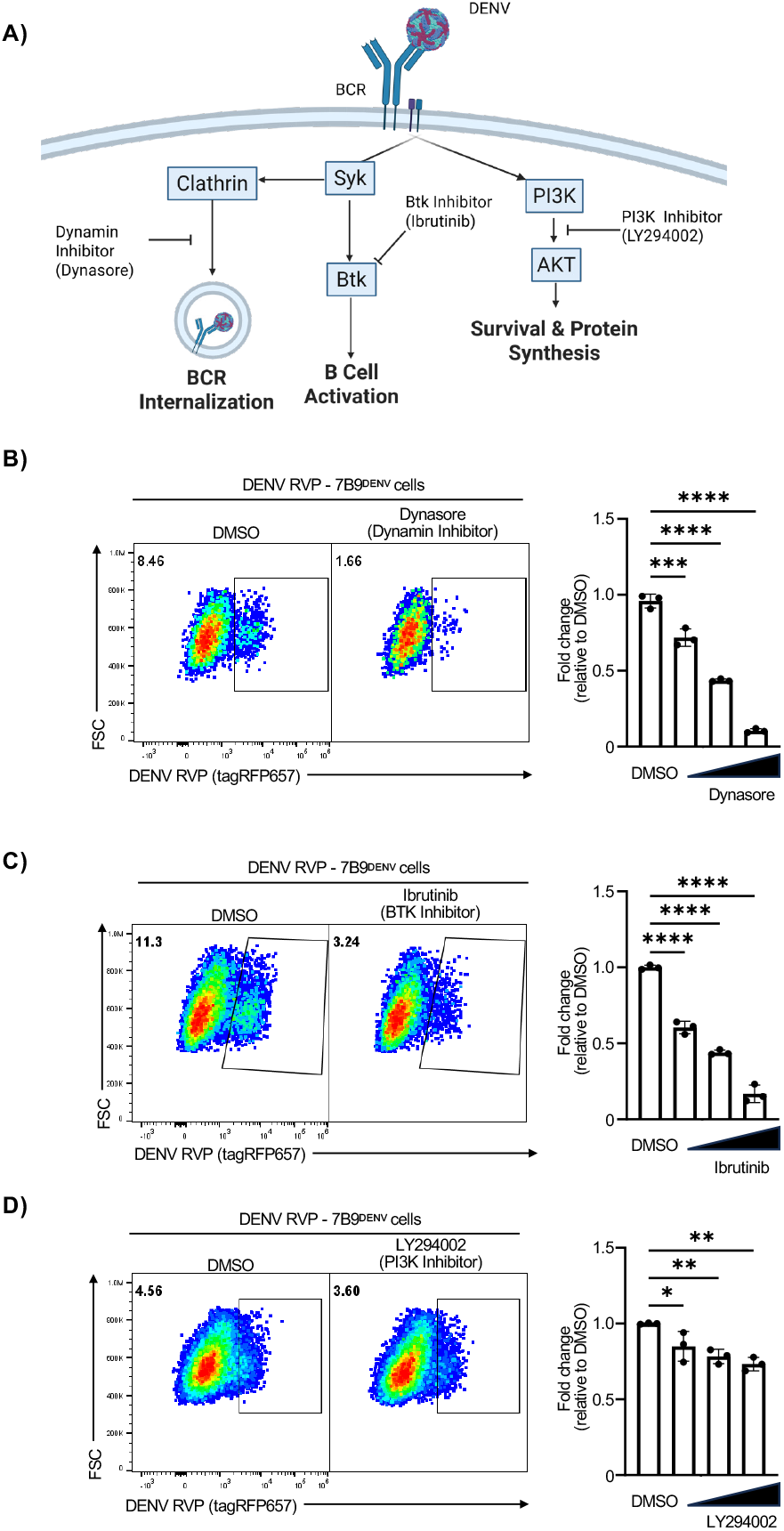
Proximal BCR signaling required for viral entry in DENV-reactive B cells. **A)** Schematic representation of BCR signaling pathways and targets of pharmacological inhibitors. Cells were pre-treated with selected inhibitor for 30 minutes prior to the addition of DENV-4 RVP. **B)** Flow cytometric analysis of DENV-4 RVP infection of 7B9^DENV^ B cells pre-treated with dynamin inhibitor Dynasore. **C)** Representative flow plots of DENV-4 RVP infection of 7B9^DENV^ B cells pre-treated with BTK inhibitor ibrutinib. **D)** Flow cytometric analysis of DENV-4 RVP infection of 7B9^DENV^ B cells pre-treated with PI3K inhibitor LY294002. Error bars +/-SEM. ^*^ *P* < 0.05, ^**^ *P* < 0.01, ^***^ *P* < 0.001, ^****^ *P* < 0.0001, One-way ANOVA with Dunnett’s multiple comparisons test, with a single pooled variance. Data are representative of two biological replicates each performed with 3 technical replicates.

Consistent with our hypothesized mechanism of action for BDE, inhibition of dynamin-mediated endocytosis with the drug Dynasore resulted in a significant and dose-dependent reduction in DENV RVP infected 7B9^DENV^ cells (**Fig. 2B**). This observation is consistent with the endocytic pathways utilized by DENV in other cell types (33, 34). Next, we sought to assess the contribution of proximal BCR-specific signaling on BDE and observed a similar reduction in the frequency of RVP infected cells upon inhibition of BTK signaling with ibrutinib (**Fig. 2C**). However, only a modest – albeit statistically significant - reduction in RVP infection was observed in cells treated with the PI3K inhibitor LY294002 (**Fig. 2D**), despite significant reduction of downstream ribosomal S6 phosphorylation (**Sup. Fig. 3A-C**). This suggests that while PI3K activity can be induced following BCR ligation by DENV, it is mostly dispensable for the initial infection of B cells via BDE. Notably, cell viability was not significantly impacted in the treated cells, with the exception of the highest concentration of Dynasore (**Sup. Fig. 3D**). To confirm the reduction in DENV RVP signal observed upon drug treatment was not due to an overall decrease in protein translation or altered cellular homeostasis, we also quantified expression of GFP within the 7B9^DENV^ cells, which is driven by a constitutive promoter contained within the retrovirus utilized to immortalize the cells. GFP expression remained stable across all conditions, indicating that the observed reductions in RVP signal resulted from decreased viral entry rather than global disruption of protein translation or cellular homeostasis (**Sup. Fig. 3E**).

### Endosomal localization of DENV in B cells

Having demonstrated the importance of BCR specificity and proximal BCR signaling to the process of BDE, we investigated the timing and subcellular trafficking of DENV following BCR engagement. We hypothesized that DENV/BCR engagement would lead to receptor internalization and subsequent localization to the late endosome, a necessary part of the DENV entry lifecycle.

7B9^DENV^ and 2F3^ZIKV^ cells were incubated with DENV-2 at 4°C for 1 hour to allow for surface binding of the virion, but not internalization. Cells were then washed, plated, temperature shifted to 37°C, and subsequently fixed at 15-, 30-, and 45-minutes post temperature shift and analyzed DENV localization and endosomes by immunofluorescence microscopy using 4G2 and LAMP-1 staining, respectively. (**Fig. 3A, 3B**). We observed a significant increase in DENV-2 and LAMP-1 colocalization over time in the DENV-specific 7B9^DENV^ cells compared to the 2F3^ZIKV^ cells (**Fig. 3B-D**). Over time, we saw diffuse DENV signal begin to form condensed puncta alongside LAMP-1 signaling inside these cells, with strong colocalization at 45 minutes post temperature shift. Despite higher levels of background signal, likely due to non-specific, low affinity binding to cells, we did not observe any significant changes in DENV-2/LAMP-1 localization in the 2F3^ZIKV^ cells (**Fig. 3B-D**). These results show that over time, DENV-2 bound to antigen-specific BCRs become internalized and subsequently localize to late endosomes.

**Figure 3.**
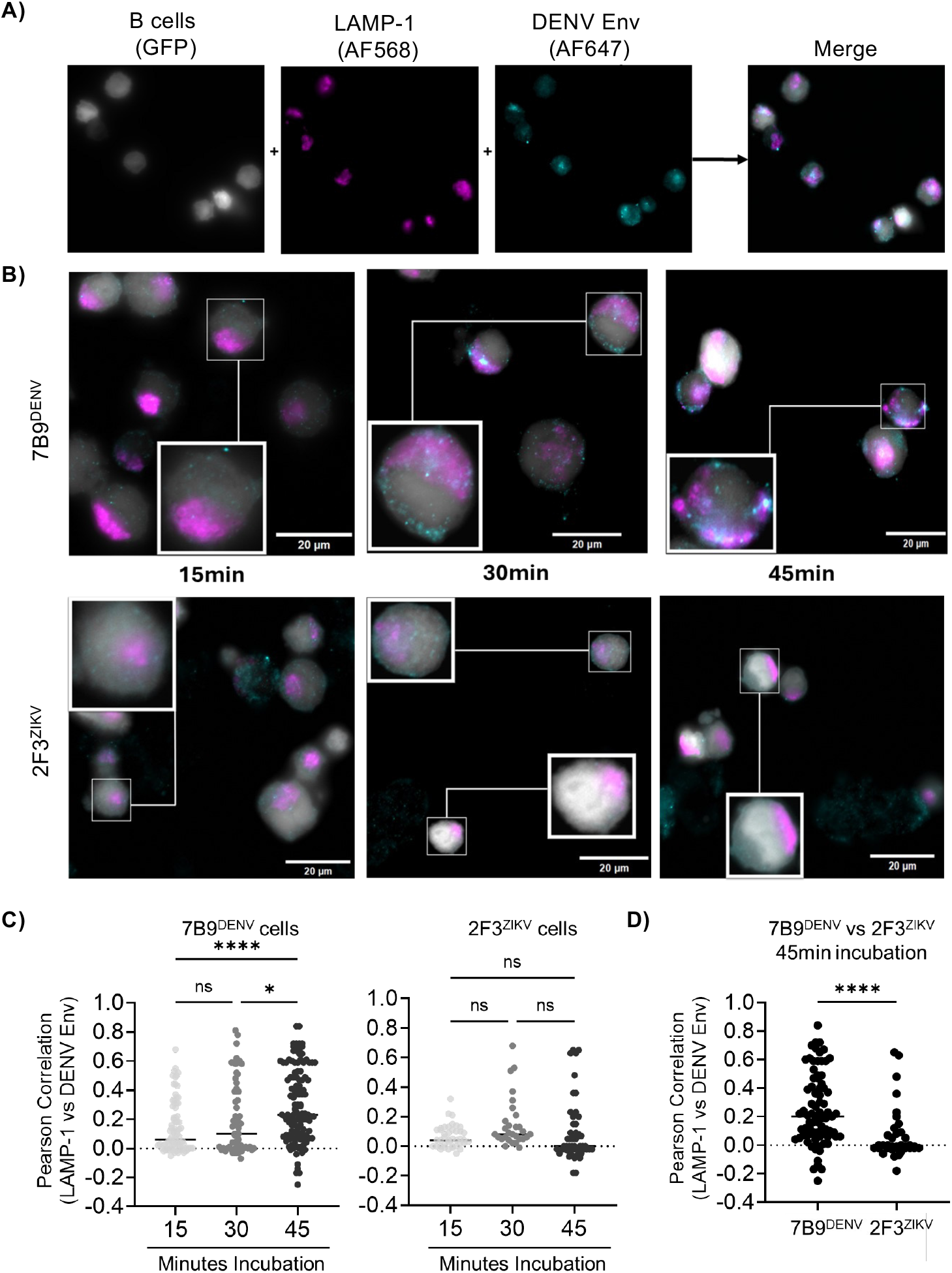
Internalization of DENV by DENV-specific B cells. **A)** Overlay fluorescent microscopy image of GFP+ B cells [grey] stained for DENV [cyan] and LAMP-1 [magenta]. **B)** DENV Internalization. Cells were infected at 4° for 2hr. Cells were fixed 15, 30, or 45min after incubation at 37° to allow for internalization, then permeabilized (1% BSA, 0.1% Triton) overnight. Cells were stained with 4G2 (mouse anti-DENV), and rabbit anti-LAMP-1 for primary staining, followed by anti-mouse (Cyan) and anti-rabbit (Magenta). **C)** Colocalization correlation image of B-cells infected through time. **D)** Colocalization correlation 45min p.i. Correlation was calculated by colocalization analysis using the Fiji image processing package for ImageJ (Coloc2). A minimum of 30 cells from each sample (n = 33 – 125) in different frames were screened, Pearson’s R value (no threshold) was scored, then plotted for grouped comparisons. Results represent pooled analysis of two independent technical replicates. ns= not significant, ^*^ *P* < 0.05, ^**^ *P* < 0.01, ^***^ *P* < 0.001. One-Way Anova with correction for multiple comparisons.

### B cells are productively infected with DENV via BCR-mediated infection

In our previous work, we demonstrated the presence of positive- and negative-strand DENV RNA in immortalized B cells expressing a DENV-reactive BCR that were exposed to DENV-2 (26). In contrast, no DENV RNA was detected in B cells expressing a ZIKV-reactive BCR following incubation with DENV-2. These results suggested that B cells may be capable of supporting active DENV replication. Additionally, since BCR signaling was involved in early infection steps and DENV was internalized in 7B9^DENV^ cells, we hypothesized that these cells may complete DENV replication, resulting in infectious virions. To test this, we exposed both 7B9^DENV^ cells and 2F3^ZIKV^ cells to DENV-2 (New Guinea C) at MOI = 10 for 24hrs, consistent with the conditions utilized in our previously published scRNAseq analysis (26). Irradiated CD40L-expressing L cell fibroblasts used as feeder cells for B-cell maintenance were exposed in a similar fashion as a negative infection control, while Vero cells were utilized as a positive control. Following DENV exposure and washing, cells were incubated to allow for viral replication and supernatants inoculated onto Vero cells to determine presence of infectious DENV by a focus forming unit assay (FFU). Strikingly, we observed robust DENV-2 production from the 7B9^DENV^ cells starting 24 hours after initial inoculation, similar in magnitude to what was observed from inoculated Vero cell cultures (**Fig. 4A, 4B**). There was also substantial infectious DENV-2 observed in 2F3^ZIKV^ cell and CD40L feeder cell cultures, likely representing residual inoculum from this high MOI experiment (**Fig. 4A, 4B**).

**Figure 4.**
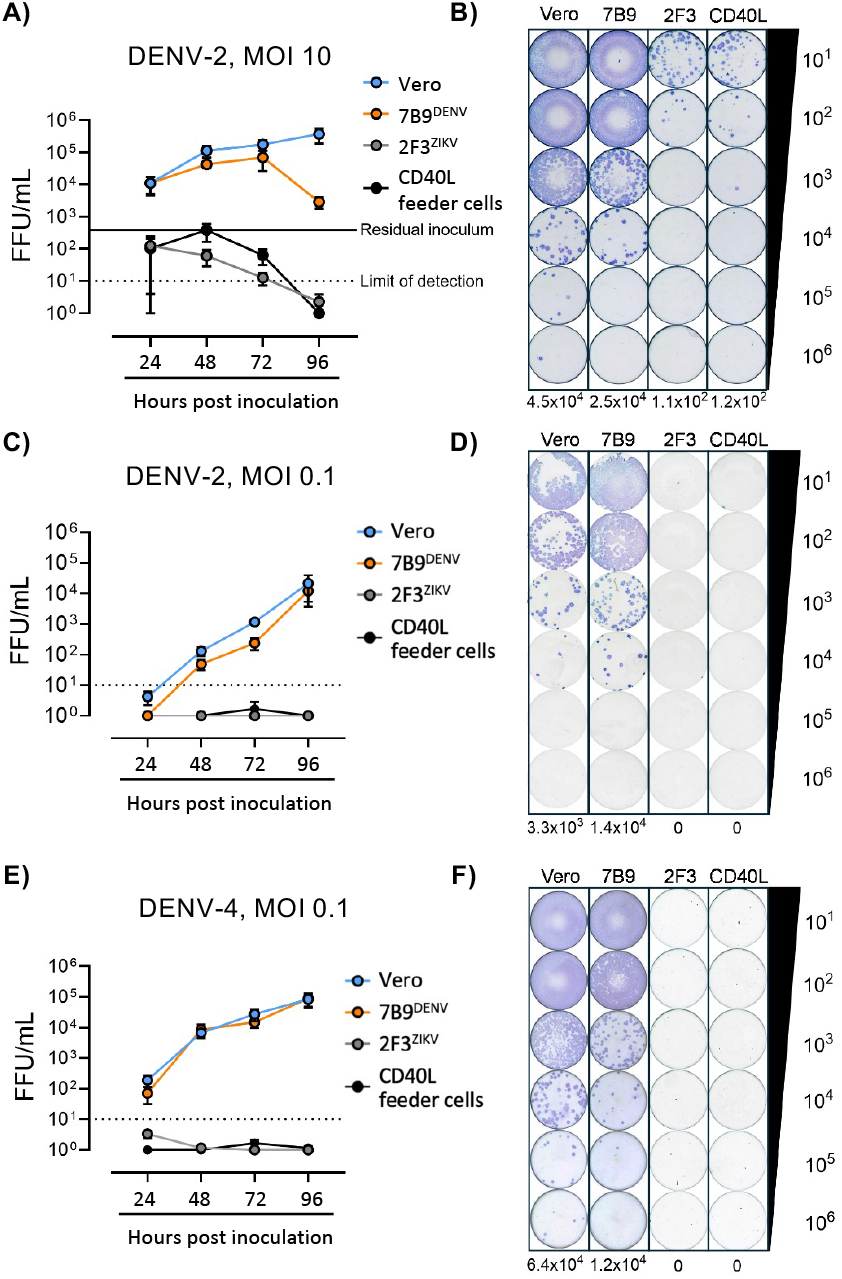
Generation of infectious DENV by DENV-specific B cells. **A)** Vero cells, 7B9^DENV^ B cells, 2F3^ZIKV^ B cells, or CD40L-expressing fibroblast feeder cells were inoculated with DENV-2/NGC at MOI = 10 for 24 hrs at 37°C, washed and cultured for the indicated times. Infectious virus titers in supernatants was determined by passage on Vero cells and assessed using a standard focus assay and data presented as focus forming units (FFU)/mL). **B)** Representative DENV-2 focus assay at MOI = 10 inoculum from 72 hr supernatants. **C)** Cells were infected with DENV-2/NGC at MOI = 0.1 for 1 hr at 37°C, washed and cultured for determination of viral titers on Vero cells in supernatants at the indicated timepoints following infection. **D)** Representative DENV-2 focus assay at MOI = 0.1 inoculum from 72 hr supernatants. **E)** Cells were infected with DENV-4/Dominica81 for 1 hr at MOI = 0.1 at 37°C followed by washing, and determination of viral titer on Vero cells at the indicated timepoints after culture. **F)** Representative DENV-4 focus assay at MOI = 0.1 inoculum from 72 hr supernatants. Experiments were performed in triplicate with two biological replicates and mean ± standard deviation is shown. Dashed lines in A, C, E indicate FFU assay limit of detection. Solid line in A indicates residual carry-through of viral inoculum.

Therefore to better characterize the production of infectious DENV from these cells, we next employed more stringent infection conditions to avoid potential cytotoxicity or cell stress associated with high viral levels and to reduce residual carry-through of viral inoculum. Accordingly, we incubated cells at an MOI of 0.1 with DENV-2 for 1 hour, followed by extensive washing and additional incubation of inoculated cultures for 24, 48, 72, and 96 hours. Under these more stringent conditions, we observed no carry-through viral inoculum (CD40L-feeder cell condition) or production of infectious DENV-2 by 2F3^ZIKV^ cells. In contrast, 7B9^DENV^ cells exhibited sustained production of infectious DENV-2 at levels similar to that observed for Vero cells inoculated under the same conditions (**Fig. 4C, 4D**). Given that the BCR of 7B9^DENV^ cells binds to DENV-4 in addition to DENV-2, we next assessed the capacity of 7B9^DENV^ cells to support replication of DENV-4. Using our stringent infectivity assay of MOI = 0.1 and 1 hr incubation with virus, we observed generation of infectious DENV-4 by 7B9^DENV^ cells to levels similar to Vero cells, while no infectious virus was produced by inoculated 2F3^ZIKV^ cells (**Fig. 4E, 4F**). These results demonstrate that DENV-specific B cells can support the replication of multiple DENV serotypes for at least 96 hours *in vitro*, producing infectious virus at similar levels to that observed from the commonly used Vero cell line.

## DISCUSSION

In this study, we provide direct evidence that dengue virus (DENV) can productively infect human B cells in a fashion dependent on B cell receptor (BCR) specificity and proximal BCR signaling. These findings expand the current paradigm of dengue immunopathogenesis, which has historically emphasized antibody-dependent enhancement (ADE) in FcγR-bearing myeloid cells, by providing mechanistic insight into how B cells themselves serve as a cellular reservoir for DENV replication. Previous studies have reported non-neutralizing DENV-specific antibodies facilitating ADE, but the role of B cells as direct targets of infection has remained underappreciated. By demonstrating productive replication and infectious particle release from DENV-specific B cells, our work provides mechanistic support for clinical observations of B cell tropism.

These results support a growing body of literature emphasizing B cells as the principal circulating cellular reservoir of DENV-infected cells (12, 19-24). Our findings work in conjunction with existing models of ADE, expanding on our understanding of DENV cellular tropism and explaining why we see such a large prevalence of DENV-infected, but non-ADE susceptible cells in circulation during acute illness. We posit that these highly susceptible B cells generated after the initial exposure aid in the establishment of DENV infection during secondary infection, where the frequency of these highly susceptible cells, paired with other risk factors such as waning neutralizing antibody titers could synergize to increase the risk of severe disease.

While this study provides additional mechanistic insight into BDE, several facets of the phenomenon require further characterization. Although we observed that proximal BCR signaling is required to establish infection, it remains unclear whether BCR signaling also influences the intrinsic permissivity of DENV-specific B cells to DENV infection. An analogous process, referred to as “intrinsic” ADE, has been described in which differential signaling through the FcR induces metabolic changes that create a more favorable environment for DENV replication. Understanding how BCR engagement impacts this process will be important, particularly given our observation that BTK inhibition decreases viral entry. Although we observed that PI3K inhibition did not affect entry, DENV can induce or dysregulate PI3K activation in other cell types (35). Given PI3K’s critical role in regulating cell survival, cellular metabolism, and protein translation, this pathway likely influences DENV replication and maturation within BDE-infected cells. Beyond these intrinsic effects, it will also be important to understand how DENV infection via BDE impacts B cell function and immunologic memory, particularly given the shift from poorly neutralizing, type-specific responses to broadly neutralizing, cross-reactive responses observed following secondary infection (36).

There are some limitations to this study that highlight the need for additional analysis. Namely, the DENV-specific human B cells utilized for this analysis were immortalized by retroviral transduction. These cells were indispensable for the analysis described in this manuscript due to the specificity of their BCR, which allowed for minimal manipulation, as well as their functional BCR signaling (26, 37). However, these immortalized IgG memory cells are maintained in a partial plasmablast-like state through persistent culturing alongside CD40L-expressing cells and IL-21 (26, 37). These in vitro cultures do not recapitulate the heterogeneity of primary human B cells, requiring further characterization in more biologically relevant models. Additionally, BCR engagement and internalization activate a complex signaling network; further study of other BCR signaling components—such as Lyn, Syk, SHP1, and Ca^2+^ flux—is required to understand the full impact of BCR signaling on DENV entry. Finally, the downstream consequences of B cell infection, including effects on antibody production, apoptosis, and immune regulation, remain to be elucidated.

In conclusion, this study has further characterized the phenomenon of BCR-dependent enhancement as a novel mechanism of viral entry for DENV. This mechanism expands our understanding of how immunologic memory shapes DENV susceptibility, and how the presence of DENV-specific memory B cells may influence an individual’s risk of infection or progression to severe disease. This study highlights the need to understand not only the contribution of antibodies as mediators of protection and pathogenesis in dengue, but how B cells themselves may contribute to risk and protection from DENV infection.

## METHODS

### Viruses & RVPs

DENV-2/NGC & DENV-4/Dominica81 stocks were prepared by propagating a low-passage inoculum (MOI = 0.5) in Vero81 cells maintained in OPTI-PRO serum free medium (Invitrogen, cat. no.12309019) supplemented with 2% fetal bovine serum (Atlanta Biologicals/R&D Systems, cat. no. S11150) and GlutaMAX (Gibco, cat. no. 35050061) at 37°C, 5% CO_2_ in a humidified incubator. Viruses were titrated by focus forming unit (FFU) assay as described (38). DENV-2 (strain Eden-2) stock was prepared by propagating a low-passage inoculum in Vero cells. Infectious particles quantified by plaque assay. DENV-4 (strain Dominica) tagRFP657 Reporter virus particles (RVP) were prepared as previously described (29). RVPs were titrated on Raji DC-SIGN+ cells, infectious units (IU) calculated based on the dilution of virus needed to infect 15-20% of cells as previously described (39).

### Cell lines

Immortalized memory B cells were previously generated by transduction of CD27^+^IgM^−^ memory B cells with a retrovirus encoding human B cell lymphoma (BCL)-6 and BCL-xL and enhanced green fluorescent protein (GFP) (26, 37). 7B9^DENV^ are an immortalized human monoclonal B-cell line expressing an IgG isotype BCR that binds, but poorly neutralizes, DENV-1 to -4 and ZIKV generated from a flavivirus-naïve volunteer who was vaccinated with DENV-1Δ30 and challenged 270 days later with DENV-2Δ30 Tonga (NCT02392325). B cells were isolated 90 days following DENV-2Δ30 challenge. 2F3^ZIKV^ are an immortalized human monoclonal B-cell line expressing a ZIKV-specific IgG isotype BCR that binds ZIKV, but not DENV and was obtained from an exposed volunteer (DT165) approximately 1 year following ZIKV infection. Cells were maintained in Iscove’s modified Dulbecco’s medium (IMDM, Invitrogen, cat. no.12200-028) supplemented with 8% fetal bovine serum (FBS) and 1% pen/strep in 24-well plates with 100,000 irradiated CD40L-expressing mouse L cell fibroblasts and 25 ng/mL rhIL-21 (Peprotech, cat. no. 200-21-1MG) as previously described (37). Raji cells overexpressing DC-SIGN+ (CD209L) (BEI, ARP-9945) were cultured in complete RPMI-1640 medium (complete RPMI) supplemented with 10% FBS and 100 U/ml penicillin–streptomycin. All cells cultured at 37°C, 5% CO_2_, and 85% humidity.

### Flow cytometry

Immortalized B cells were stained for viability (L/D) with Fixable Viability Dye eFluor 780 (Invitrogen, cat. no. 65-0865-14) according to the manufacturer’s protocol and washed with FACS buffer (PBS + 1% BSA). 100 ng of AviTagged and biotinylated DENV-2 or DENV-4 recombinant E protein stabilized dimers (DV2 SC.14 and DV4 SC, respectively, a kind gift of Brian Kuhlman and Aravinda DeSilva, University of North Carolina, Chapel Hill, (28)) were labeled with 0.5 μL of streptavidin-PE (Invitrogen, cat. no. 12-4317-87) in a volume of ∼8 μL for 30 minutes on ice. 100,000 B cells were then stained with PE-labeled recE on ice for 30 min. Following a wash with FACS buffer and final staining with anti-human IgG-APC (clone: M1310G05, BioLegend, cat. no. 410712) for 30min. Events were captured on a Beckman Cytoflex at the Microscopy Imaging and Cytometry at the University of Vermont (RRID# SCR_018821) and analyzed with FlowJo version 10.

DENV-4 RVP infection assays were collected on a ThermoFisher Attune NXT flow cytometer, detecting 7B9^DENV^ or 2F3^ZIKV^ cells through detection of GFP. RVP infection was detected through the expression of tagRFP657. Data collection was performed on a ThermoFisher Attune NXT flow cytometer and analyzed using FlowJo v10.2 (Becton Dickinson).

### DENV-4 tagRFP657 RVP Infection of human B cells

7B9^DENV^ or 2F3^ZIKV^ cells were plated at 5×10^4^ cells per well. DENV-4 RVP was titrated in two-fold serial dilutions. Cells were incubated for 4 hours at 37°C, followed by wash and trypsinization to remove remaining surface bound RVP. Then cells were incubated an additional 20 hours at 37°C. Cells were washed twice with PBS + 2% FBS then fixed using 4% PFA and detected by flow cytometry. MOI calculation for titration was back-calculated based on IU calculations on Raji DC-SIGNR cells.

### DENV-2 recombinant E Dimer infection blockade

DENV-2 recombinant Envelope protein dimers (recE) were received under MTA from Aravinda M. de Silva at University of North Carolina, Chapel Hill. 7B9^DENV^ or Raji DC-SIGN+ cells were plated at 5×10^4^ cells per well and chilled to 4°C before no treatment or addition of 0.1μg/mL or 1.0μg/mL of the DENV-2 recE. Cells were incubated at 4°C for 30 minutes, followed by the addition of DENV-4 RVP at an MOI of 10. Cells were incubated an additional 30 minutes at 4°C before three washes using cell culture medium. Cells were resuspended in fresh cell culture media and incubated for 24 hours at 37°C. Cells were washed twice with PBS + 2% FBS before fixation using 4% PFA and analyzed by flow cytometry.

### Inhibition of viral entry by pharmacological inhibitor treatment

7B9^DENV^ cells were plated at 5×10^4^ cells per well and pre-treated for 30 minutes at 37°C with DMSO or various concentrations of pharmacological inhibitors: 12.5μM, 50μM, and 200μM of dynamin inhibitor Dynasore (Abcam, cat. no. ab120192); 1.56μM, 12.5μM, and 50μM of BTK inhibitor Ibrutinib (Selleck, cat. no. S2680); or 1.56μM, 12.5μM, and 50μM of PI3K inhibitor LY294002 (Selleck, cat. no. S1105). Following pre-treatment, DENV-4 RVP was added at an MOI of 2.5 and incubated for 4 hours in the presence of drug. Cells were then washed twice using cell culture medium and trypsinized then washed again to remove surface bound RVP before resuspension in fresh cell culture media and incubated at 37°C for 20 hours. Cells were then washed twice with PBS + 2% FBS before fixation using 4% PFA and analyzed by flow cytometry.

Inhibitor treatment effect on cell viability in the absence of RVP was performed under the same conditions as described above then assessed by metabolic conversion of MTT tetrazolium into formazan utilizing the CellTiter 96® Non-Radioactive Cell Proliferation Assay (MTT) (Promega, cat. no. G4000). 1% Triton-X (Thermofisher, cat. no. A16046.AE) was used as a positive control for cell death. Data was collected on a Biotek µQuant plate reader at 570 nm.

Pharmacological inhibition of PI3K was assessed via flow cytometry through the detection of phosphorylated ribosomal protein S6 at Ser235 and Ser236 as a direct downstream measurement of activated PI3K. Cells were treated with DMSO (Thermofisher, cat. no. 036480.AP) or 10μM LY294002 for 30 minutes. Then washed twice with PBS + 2% FBS. Then fixed using 4% PFA and stained utilizing AF647 labelled phospho-S6 monoclonal cupk43k (Thermofisher, cat. no. 740057MP647) diluted 1:1000 in PBS + 2% BSA. Data collection was performed on a ThermoFisher Attune NXT flow cytometer and analyzed using FlowJo v10.2 (Becton Dickinson).

### Detection of DENV internalization and endosomal localization by microscopy

Ibidi 8-well chamber slides (Ibidi, cat. no. 80826-120) were coated with Histogrip (ThermoFisher, cat. no. 008050) to promote cell adhesion to slide. 7B9^DENV^ or 2F3^ZIKV^ cells were chilled to 4°C prior to the addition of DENV-2 EDEN live virus at MOI of 10. Cells were incubated for 1 hour at 4°C to allow surface binding before washing. Cells were then plated at 1×10^5^ per well and allowed to adhere at 4°C for an additional 1 hour. Cells were then temperature switched and incubated for 15, 30, and 45 minutes at 37°C then fixed using 4% PFA. Cells were then permeabilized using PBS supplemented with 0.1% Triton-X and 1% BSA. Cells were stained overnight at 4°C with primary antibodies diluted in perm buffer: mouse cross-reactive anti-flavivirus envelope (E) protein monoclonal antibody 4G2 diluted to 4ug/mL (Envigo, cat. no. 80216718); and rabbit anti-human LAMP1 monoclonal antibody D2D11 diluted 1:100 (Cell Signaling, cat. no. 9091). Followed by two washes in perm buffer before secondary staining in perm buffer, each diluted 1:1000: donkey anti-mouse IgG AF647 (Jackson ImmunoResearch Inc, cat. no. 715-605-151); donkey anti-rabbit IgG AF568 (ThermoFisher, cat. no. A10037). After staining, cells were washed twice in PBS. Intracellular GFP was utilized to detect immortalized B cells and used as cytoplasm dye. Images were collected on a Marianas microscopy system (3i) enclosed in an environmental chamber (Okolab) consisting of a Zeiss Axio Observer 7 equipped with a X-Cite mini + light source (Excelitas) and a Prime BSI Express CMOS camera (Photometrics). Images were taken at 63× magnification, with each image created using SlideBook 6 (3i) before exporting to Fiji for analysis. Individual cells were then analyzed for colocalization of LAMP1 and DENV by Pearson Coefficient utilizing Coloc 2 function in Fiji.

### Quantification of infectious DENV

Immortalized B cells (250,000 cells) were cultured in complete IMDM in 24-well plates on CD40L cell (100,000 cells) monolayer and supplemented with rhIL-21. Irradiated CD40L-expressing L cells (350,000 cells, to account for total cell number in immortalized B cell conditions) cultured in complete IMDM, and Vero cells (350,000 cells) cultured in OPTI-MEM 2% FBS and GlutaMAX, were included as negative and positive controls respectively. Cells were exposed to DENV at MOIs of 10 or 0.1 at 37°C for 24 hours or 1 hour as described in figure legends. Infected cells were then washed three times with DPBS (Invitrogen, cat. no. 14190-144) and cultured in complete IMDM+ rhIL-21 at 37°C for an additional 24, 48, 72, or 96 hours. Supernatants were collected at each timepoint and frozen at -80°C until further use. Supernatants were then diluted in a range of 10-fold dilutions (neat, 10^−1^ to 10^−7^) in virus titration diluent (Opti-MEM-no glutamax (Invitrogen cat. no. 31985-070), 2% heat inactivated FBS, and 10mL L-glutamine 200mM). 100 μL of each dilution were applied to confluent Vero cell monolayers in 24 well plates for 1 hour at 37°C, followed by addition of 1 mL of overlay media (Opti-MEM + Glutamax (Gibco, cat. no. 51985.091), 5g Methylcellulose, 2% heat inactivated FBS, and 0.5 mL Gentamicin 50 mg/mL (Gibco, cat. no. 15750.060)) and incubation for 4 days at 37°C. Next, overlay medium was removed by flicking and tapping plates, and monolayers were washed twice with 1 X PBS, and fixed with 50%/50% Methanol/Acetone for 40 min at 4°C, followed by blocking for 10 min with Antibody Dilution Buffer (5% Oxoid non-fat dry milk in 1× PBS), washed again twice with PBS, and incubated with 4G2 (0.8 ng/mL, HB112 hybridoma from ATCC) in blocking buffer, followed by incubation at 37°C for 1 hour. Plates were then washed twice with PBS and incubated with HRP-labeled secondary goat anti-mouse antibody (1:2000, KPL, cat. no. 074-1806) at 37°C for 1 hour, washed with PBS, air-dried for 5 minutes, and treated with 160 µL TrueBlue substrate (SeraCare, cat. no. 5510-0030), allowing foci to develop for ∼15 minutes. The developed plaques were then counted by eye, and virus titers calculated as previously described (38).

## Statistical Analysis

All statistical analyses were performed using GraphPad Prism Software (GraphPad Software, La Jolla, CA, USA). A P value <.05 was considered significant. ANOVA was used for the comparison of multiple conditions.

## Acknowledgements

We gratefully acknowledge excellent technical assistance provided by S M Jamil Mahmud of the SUNY Upstate Medical University Flow Cytometry Core, as well as Elizabeth Kurtz and Nathan Roy for their assistance with microscopy. We also wish to thank Ted Pierson and Stephen Whitehead for the constructs and protocols for reporter virus production. We are grateful to Aravinda M. de Silva and Brian Kuhlman for providing the recombinant DENV-2 and DENV-4 E dimers. 4G2 mAb was received under MTA from Walter Reed Army Institute of Research (WRAIR). The following reagent was obtained through the NIH HIV Reagent Program, Division of AIDS, NIAID, NIH: Raji DC-SIGN+ Cells, ARP-9945, contributed by Dr. Li Wu and Dr. Vineet N. KewalRamani. Vero81 cells were a kind gift from Steve Whitehead (NIAID).

## Data availability

The authors confirm that the data supporting the findings of this study are available and included within this article.

## Funding

Funding for this research was provided by the National Institute of Allergy and Infectious Diseases (NIAID) R01AI192874 (A.T.W and S.A.D.) and U19AI181960 (S.A.D.) and by the National Institute of General Medical Sciences P20GM125498 (S.A.D.). The funders had no role in study design, data collection and analysis, decision to publish, or preparation of the manuscript.

